# DREM: integrating resistance and domain probabilities to prioritize plant immune receptors

**DOI:** 10.64898/2026.10.21.760626

**Authors:** Rita Dublino, Vincenzo Carandente, Giancarlo Sperlì, Mara Ercolano

**Author notes:** Corresponding author: Mara Ercolano.

## Abstract

Plant resistance (R) genes, key players in the immune system, have traditionally been identified with HMM-based tools relying on sequence homology, whose sensitivity decreases for divergent or non-canonical resistance proteins. Here, DREM (Deep Resistance gene Evaluator Model) is presented as a deep learning framework that learns the compositional properties of resistance proteins from sequence alone rather than searching for predefined motifs. The framework combines two bidirectional long short-term memory networks with attention: a binary resistance classifier and a multi-label classifier of resistance domains. Validation on experimentally characterized proteins from RefPlantNLR recovered 99.8% of sequences, and on 15,000 held-out proteins DREM outperformed the deep learning predictor PRGminer (Matthews correlation coefficient 0.938 vs 0.820), with about two-thirds fewer classification errors. Across five plant proteomes, DREM identified 4,046 High Confidence or Candidate proteins; canonical domain architecture was confirmed in 75.5%, whereas 992 carried strong DREM signals without detectable Pfam domains, and 54 of 55 characterized nucleotide-binding leucine-rich repeat (NLR) loci ranked within the top 1.7% of their proteomes. Unlike domain-gated pipelines such as R-Predictor, DREM also recovered experimentally validated atypical NLRs, RPW8 proteins and non-receptor resistance genes. Attention did not concentrate on canonical ATP-binding motifs, consistent with the model drawing on sequence information beyond conserved domain signatures. Lightweight and applicable to complete proteomes, DREM complements domain-based annotation with a calibrated ranking. It links genome-wide prediction to the selection of resistance genes for functional validation, including atypical receptors missed by profile-based tools.

## 1. INTRODUCTION

Plant resistance genes play a central role in plant immunity and represent an important target for crop breeding (Jones *et al.,* 2024). Over the last thirty years, scientists have characterized NLRs (nucleotide-binding leucine-rich repeat proteins), RLKs (receptor-like kinases), and RLPs (receptor-like proteins) and learned how plants recognize pathogens and activate defense (Ngou *et al.,* 2022). However, the strong selective pressure imposed by pathogen evolution makes R-genes among the most diverse and rapidly changing sequences of the plant genome (Ercolano *et al.,* 2012; Ercolano *et al.,* 2022; Ngou *et al.,* 2024).

Identifying resistance genes is a critical issue: it clarifies how plants evolve to fight pathogens, and it gives plant breeders tools to develop better crops, especially as climate change drives new disease threats. Discovery of plant resistance genes from genomic sequences remains challenging due to their sequence diversity, multidomain architecture, and complex evolutionary history. Traditionally, the discovery of R-genes was based almost exclusively on sequence homology, with HMMs (Hidden Markov Models) serving as the standard (Osuna-Cruz *et al.,* 2018; Paysan-Lafosse *et al.,* 2025). Several bioinformatic tools, such as NLR-Parser, RGAugury, and DRAGO2, employ predefined motifs and domain signatures to classify sequences (Steuernagel *et al.,* 2015; Li *et al.,* 2016; Calle García *et al.,* 2022). Further implementations like NLRtracker and similar pipelines integrate multiple detection methods with curated reference databases such as RefPlantNLR or PRGdb to assign functional domains and classify genes (Kourelis *et al.,* 2021; Calle García *et al*., 2022). More recently, machine-learning predictors have been proposed, including PRGminer, a deep neural network that classifies proteins as R-genes and assigns them to eight classes (Duhan and Kaundal, 2025), and R-Predictor, a pipeline combining protein language models with domain-based modules (Liu et al., 2026). R-Predictor assigns proteins to domain topologies and therefore relies on detectable Pfam domains, whereas PRGminer returns class probabilities without an explicit measure of structural coherence. The fundamental limitation of this pattern-matching strategy is that tools can only identify what is already known (Sapoval *et al.,* 2022). While highly effective for conserved sequences, these methods lose sensitivity when proteins have undergone substantial evolutionary divergence, lost characteristic domains, or acquired atypical structural organizations (Hofberger *et al.,* 2014). This creates a systematic blind spot: resistance genes with divergent sequences or unusual domain combinations remain undetected, limiting discovery of the full genetic diversity available for breeding and functional characterization.

Deep learning offers an alternative paradigm: moving from recognizing predefined patterns to understanding the functional compositional logic of a protein directly from data (Senior *et al.,* 2019; Boadu *et al.,* 2025). To capture this complexity, recurrent neural network architecture, such as BiLSTM (Bidirectional Long Short-Term Memory) model protein sequences as ordered series of residues. Unlike linear models, a BiLSTM is designed to process sequential data where position and context are decisive (Wang *et al.,* 2022). The bidirectional nature of this network allows it to process protein sequences from the N-terminus to the C-terminus and vice versa simultaneously. This approach enables the model to capture structural dependencies that local window-based methods or standard profiles often ignore (Liu *et al.,* 2022). While an HMM looks for a specific match to a signature, the BiLSTM learns a latent representation of the protein. It identifies statistical regularities in amino acid distribution that define a function, even when specific motifs are missing.

Furthermore, attention mechanisms have emerged as a promising implementation for performing sequential scanning. These mechanisms improve model performance, allowing the network to focus on the most informative regions of a sequence (Ranjan *et al.,* 2023; Ke *et al.,* 2024). Attention weights make it possible to visualize which part of the sequence, often corresponding to signaling interfaces or recognition domains, drive the model’s decision (Liu and Gong, 2019). It provides a level of biological interpretability that is essential for validating new candidate genes.

In this paper, DREM (Deep Resistance gene Evaluator Model) is presented as a framework that implements a BiLSTM strategy to identify resistance genes based on a double assessment: one for binary classification (resistance vs. non-resistance) and one for the multi-label prediction of key structural domains (NBS, LRR, TIR, CC, and kinase). DREM was trained on a balanced subset derived from a comprehensive collection of nearly one million sequences. Two complementary metrics were designed to evaluate its performance: The Structural Confidence Score (SCS) and the Protein Resistance Score (PRS), which evaluate both sequence-level resistance signatures and the domain consistency of DREM predictions. DREM was compared with PRGminer and with the domain-gating steps of R-Predictor, and its ranking was evaluated on characterized NLR loci in complete proteomes. Its predictive power was further tested on independent, highly curated datasets and the complete proteomes of five plant species of major agricultural and scientific importance: *Arabidopsis thaliana, Oryza sativa*, *Solanum lycopersicum*, *Vitis vinifera* and *Cucurbita pepo*.

The goal was not just to introduce a new analysis tool, but rather to define a framework for integrating deep learning into comparative genomics. DREM explores the dark matter of plant immunity, transforming the ability to map and utilize the genetic diversity essential for the future of agriculture.

## 2. RESULTS

### 2.1. Convergence analysis and training configuration

DREM processes raw amino acid sequences through two networks of identical architecture (embedding, two-layer BiLSTM and additive attention pooling), trained separately as a binary resistance classifier and a multi-label domain classifier, whose outputs are combined into a composite confidence score (Fig. 1).

**Fig. 1.**
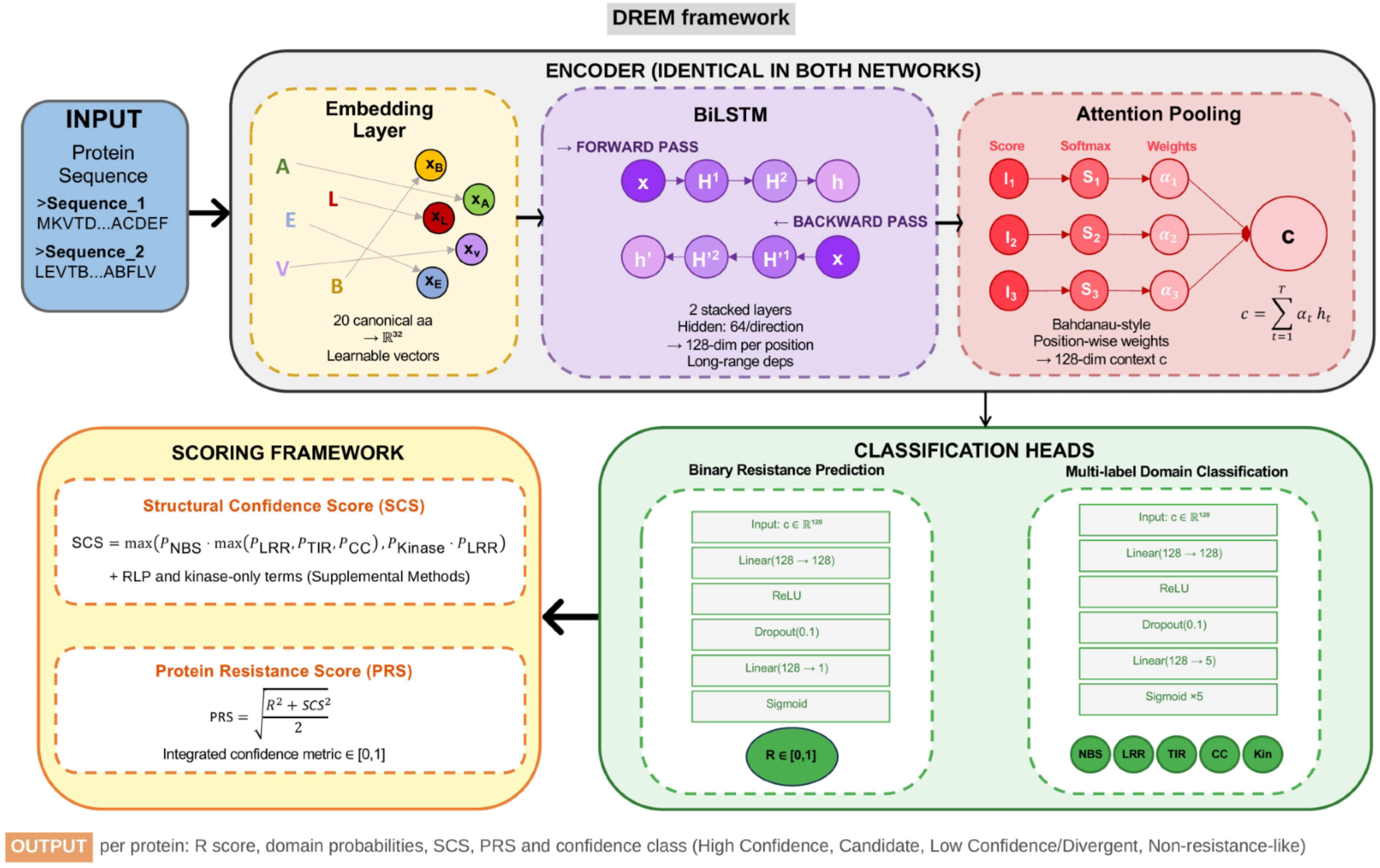
Architectural workflow of the DREM framework. Schematic representation showing the DREM processing stages: input and embedding layer for amino acid sequences; an encoder, identical in both networks, featuring a two-layer BiLSTM and Bahdanau-style attention pooling; classification heads for binary resistance prediction and multi-label domain classification; and the scoring framework integrating both outputs into the Structural Confidence Score and the final Protein Resistance Score. All layer dimensions are indicated within the corresponding panels.

Convergence was systematically evaluated across three dataset sizes: 50,000, 100,000, and 150,000 sequences (hereafter DS50, DS100, and DS150) and multiple epoch configurations (10-50 epochs) (Methods; Table S1).

The binary resistance classifier displayed dataset-dependent convergence behavior. In DS50 convergence peaked at epoch 6 with notable oscillations reflecting sampling variability at smaller scale while in DS100 convergence stabilized by epoch 8. In DS150 validation loss decreased monotonically through epoch 10 (Val loss: 0.0579), indicating the model had not fully saturated its representational capacity, yet downstream test performance was already high (ROC-AUC: 0.9967, MCC: 0.9383). The multi-label domain classifier exhibited a consistent pattern independent of max-epoch setting: minimum validation loss was invariably reached at epoch 9, after which validation loss increased monotonically despite continued training loss decrease, providing clear evidence of overfitting beyond epoch 9. Model checkpointing at this optimal epoch ensured identical test performance across all epoch configurations, confirming that extended training provided no overall benefit. Both classifiers were therefore trained on DS150 for 10 epochs for all downstream analyses. This configuration balanced performance and computational efficiency while achieving lowest validation loss across all tested conditions. To assess its stability, this configuration was evaluated using 5-fold stratified cross-validation. In 5-fold cross-validation on 150,000 sequences, the binary resistance classifier achieved 97.11% ± 0.12% accuracy (F1-score = 97.12% ± 0.11%, AUC = 99.70% ± 0.03%), and the multi-label domain classifier achieved 93.80% ± 0.13% accuracy (F1-score = 94.45% ± 0.06%, AUC = 99.63% ± 0.03%) (Table S2). The very low standard deviations across folds indicate stable performance across data partitions.

### 2.2. Binary resistance classification performance

The binary resistance classifier was evaluated on the held-out test set (15,000 sequences from the DS150), assessing its ability to discriminate between resistance and non-resistance protein sequences from raw amino acid input alone. Performance was assessed using accuracy, ROC-AUC, Matthews Correlation Coefficient (MCC), F1-score, precision, and recall, all computed at the default decision threshold of 0.5.

On the DS150, the binary classifier achieved high discriminative performance. Accuracy reached 0.9691, ROC-AUC attained 0.9967, and MCC reached 0.9383 (Fig. 2A; Table S1). F1-score was 0.9691, with precision of 0.9743 and recall of 0.9641. Per-class analysis revealed both resistance (precision 0.974, recall 0.964) and non-resistance (precision 0.964, recall 0.974) classes predicted with comparable metrics, demonstrating that the balanced training strategy prevented the model from developing a systematic class preference.

**Fig. 2.**
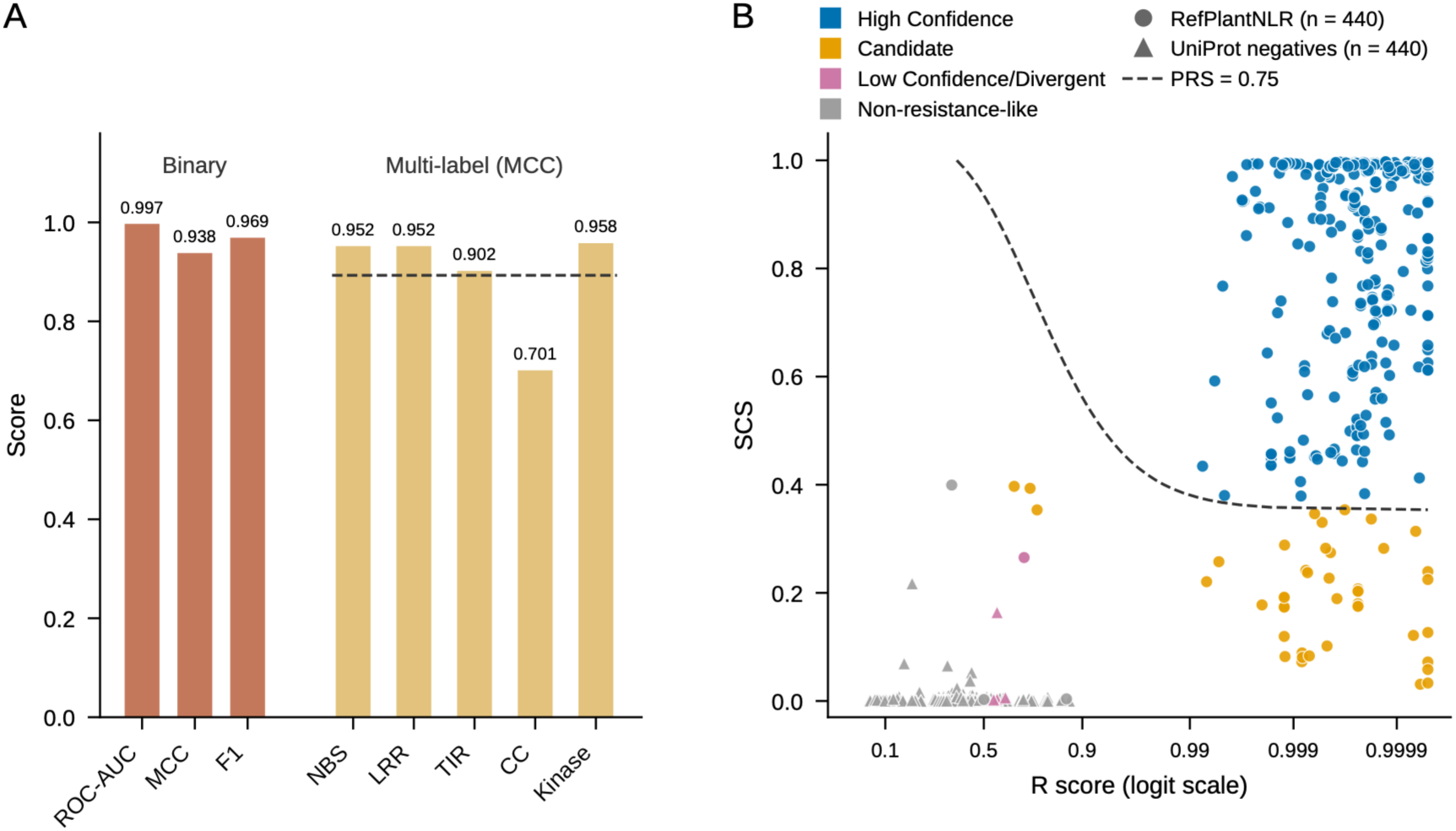
Classification performance and external validation of DREM. A: Performance of the binary classifier (ROC-AUC, MCC and F1) and per-domain MCC of the multi-label classifier on the DS150 held-out test set (n = 15,000); the dashed line indicates the macro-averaged MCC (0.893). B: R score and structural confidence score of RefPlantNLR proteins (circles) and UniProt non-resistance proteins (triangles), colored by confidence class; the dashed line marks PRS = 0.75. MCC, Matthews correlation coefficient; SCS, Structural Confidence Score; PRS, Protein Resistance Score.

The ROC-AUC of 0.9967 demonstrates well-separated probability distributions across the full decision threshold range. This balance is essential for proteome-wide inference, where asymmetric predictions could either saturate the candidate set with spurious predictions or miss genuine resistance genes.

Performance improved consistently with training set size (Table S1). ROC-AUC improved from 0.9896 (DS50) to 0.9953 (DS100) and 0.9967 (DS150), with diminishing increments (+0.0057 and +0.0014) that reflect the compression of this metric near its upper bound. MCC and accuracy, which are more sensitive in this range, continued to improve at a steady rate across the same interval (MCC: 0.9222, 0.9294, 0.9383). DS150 was therefore selected as the final configuration: performance had not yet plateaued, but was already sufficient for proteome-wide application, and further expansion of the training set would have increased computational cost without a corresponding gain in operational utility. The continuous output of the binary classifier, designated the Resistance Score (R score), represents the model’s confidence that a given sequence encodes a resistance protein. This score (range [0,1]) constitutes the primary output of DREM’s first classification step and R ≥ 0.5 is used as the decision threshold of the binary classifier, whereas candidate prioritization relies on the composite scores described below.

### 2.3. Multi-label domain classification performance

The multi-label domain classifier was evaluated on the held-out test set (15,000 sequences from the DS150), simultaneously predicting five canonical structural domains: NBS, LRR, TIR, CC, and Kinase. At the global multi-label level, Exact Match Accuracy reached 0.9367, indicating that all five domain labels were simultaneously correct in over 93% of test sequences. Hamming Loss was 0.0136 (1.4% average per-label error rate), indicating coherent joint predictions. Aggregate metrics showed Micro F1 of 0.95, Macro F1 of 0.90, macro-averaged ROC-AUC of 0.992, and PR-AUC of 0.940 (Table S1).

Per-label performance varied in biologically interpretable patterns (Fig. 2A; Table S3). NBS and LRR domains achieved the highest ROC-AUC (0.999 for both; MCC: 0.952), reflecting their strong evolutionary conservation at the sequence level. Kinase domains demonstrated high performance (ROC-AUC: 0.998, MCC: 0.958, F1: 0.972). TIR achieved intermediate but strong performance (ROC-AUC: 0.990, MCC: 0.902, F1: 0.903), while CC domain prediction showed distinct performance characteristics (ROC-AUC: 0.976, MCC: 0.701, F1: 0.703) reflecting the intrinsic sequence heterogeneity of coiled-coil regions, which are among the least conserved structural elements at the primary sequence level.

Performance improved with dataset size (Table S1): macro-averaged ROC-AUC increased from 0.987 (DS50) to 0.992 (DS100 and DS150), and macro MCC from 0.854 to 0.893. This scaling reflects the benefit of increased training diversity.

### 2.4. Biological validation on known resistance proteins

To assess whether DREM is able to infer resistance proteins with direct experimental support, two independent validation analyses were performed on sequences deliberately excluded from training to prevent data leakage and ensure unbiased assessment.

The first set consisted of 152 curated PRGdb resistance proteins with direct literature support. After sequence-level deduplication, 148 non-redundant proteins were retained for biological validation. The second set consisted of 440 resistance proteins from RefPlantNLR, an independently curated database of plant NLR proteins. Together, these datasets provided 588 experimentally supported resistance proteins for external positive validation. As a reference, on the DS150 held-out test set (15,000 proteins; see above), the mean R score was 0.9591 for resistance proteins and 0.0307 for non-resistance proteins, with median values of 0.9998 and 0.0001, respectively, confirming a strong separation between the two classes (Data Set S1).

On the independent PRGdb positive validation set, DREM assigned R ≥ 0.5 to 138 out of 148 proteins, corresponding to a recovery rate of 0.9324 (Data Set S2). Among these, 127 proteins reached the high-confidence threshold R ≥ 0.8, corresponding to 0.8581 of the validation set.

The mean R score was 0.9278, while the median R score was 0.9996, with an interquartile range from 0.9984 to 0.9998. This distribution indicates that most curated PRGdb resistance proteins received near-maximal resistance scores, although a small subset showed lower scores and accounted for the reduced mean relative to the median. The RefPlantNLR validation set showed even stronger recovery. DREM assigned R score ≥ 0.5 to 439 out of 440 proteins, corresponding to a recovery rate of 0.9977 (Data Set S3). High-confidence predictions with R score ≥ 0.8 were obtained for 433 proteins, corresponding to 0.9841 of the dataset. In addition, 388 proteins showed PRS ≥ 0.75, indicating that 0.8818 of RefPlantNLR proteins were not only recovered as resistance-like by the binary classifier, but also received high composite prioritization scores integrating R score and SCS (Fig. 2B). The mean R score was 0.9934 and the median was 0.9998, with an interquartile range of 0.9996 to 0.9999.

The PRGdb validation set showed a broader lower tail, consistent with the presence of curated resistance proteins with atypical, truncated, divergent or less canonical sequence features. By contrast, RefPlantNLR proteins showed a more compact high-score distribution, consistent with the predominance of canonical NLR architectures in this dataset. Overall, these results show that DREM maintains high discriminative performance on the internal test set and robustly recovers experimentally supported resistance proteins excluded from training. The concordant recovery of PRGdb and RefPlantNLR proteins supports the generalization capacity of the binary resistance classifier, while the lower-scoring PRGdb subset highlights biologically meaningful heterogeneity among validated resistance genes rather than uniform score inflation.

Per-protein predictions for the RefPlantNLR proteins and for 440 UniProt non-resistance proteins used as negative controls are provided in Data Sets S3 and S4. None of the negative controls was classified as High Confidence or Candidate, although 99 had R ≥ 0.5, confirming that the composite classification filters proteins that the R score alone would retain.

### 2.5. Proteome-wide inference

To demonstrate the scalability and overall capacity of the DREM framework, proteome-wide inference was performed on the complete proteomes of five plant species of major agricultural and scientific relevance: *Vitis vinifera* (grapevine), *Solanum lycopersicum* (tomato), *Oryza sativa* (rice), *Arabidopsis thaliana*, and *Cucurbita pepo* (squash). These species were selected to represent broad taxonomic diversity across monocots and dicots and to include crops of primary economic and food security importance. Complete proteome sequences were retrieved from their respective Ensembl Plants repositories and processed without species-specific adaptation or retraining, applying the identical encoding, truncation, and classification pipeline described in Methods. The total number of protein sequences analyzed ranged from 27,862 (*C. pepo*) to 48,306 (*A. thaliana*) reflecting differences in proteome annotation completeness across species and genomic reference quality.

Proteins were assigned to four confidence classes by rules combining R and domain probabilities, with PRS separating High Confidence from Candidate proteins (Methods). The proportion of proteins classified as High Confidence or Candidate, collectively representing the putative resistance gene complement predicted by DREM, ranged from 1.42% of the total proteome in *C. pepo* (397 out of 27,862 proteins) to 2.93% in *V. vinifera* (1,205 out of 41,097 proteins), with intermediate values in the other species (Table 1). These proportions are broadly consistent with published estimates of resistance gene density in plant genomes, which typically reflect differences in genome annotation quality, evolutionary history, and lineage-specific NLR expansions depending on species and annotation (Li *et al.,* 2023). *Vitis vinifera* yielded the highest absolute number of High Confidence predictions (881), *Arabidopsis thaliana* followed with 802 High Confidence proteins, *Oryza sativa* (506), *Solanum lycopersicum* (378), and *Cucurbita pepo* (275) showed progressively lower High Confidence counts. Candidate proteins meeting the primary structural criteria (well-predicted domains), but with PRS ≤ 0.75, were identified in all five species, ranging from 122 (*C. pepo*) to 324 (*V. vinifera*). These represent a lower-confidence set of predictions warranting further experimental investigation, as they exhibit strong domain signals but lower composite scores, potentially indicating borderline cases, divergent homologs, or genuine resistance genes with atypical sequence features.

**Table 1.** Proteome-wide DREM classification and HMMER validation in five plant species. Values are n (%). Proteins were assigned to four confidence classes with the rules described in Methods; percentages of the four classes are relative to the total number of proteins in each species. HMMER+ proteins carry at least one Pfam domain hit (E-value < 1 × 10⁻⁵); their percentage is relative to High Confidence plus Candidate proteins. HMMER− proteins have no detectable canonical domain. HC, High Confidence; Cand., Candidate; LC/D, Low Confidence/Divergent; NRL, Non-resistance-like.

| Species | Proteins | HC | Cand. | LC/D | NRL | HMMER+ | HMMER– |
| --- | --- | --- | --- | --- | --- | --- | --- |
| *Arabidopsis thaliana* | 48,306 | 802 (1.66) | 280 (0.58) | 4,155 (8.60) | 43,069 (89.16) | 811 (74.95) | 271 |
| *Oryza sativa* | 42,579 | 506 (1.19) | 244 (0.57) | 3,257 (7.65) | 38,572 (90.59) | 585 (78.00) | 165 |
| *Solanum lycopersicum* | 34,361 | 378 (1.10) | 234 (0.68) | 2,687 (7.82) | 31,062 (90.40) | 450 (73.53) | 162 |
| *Vitis vinifera* | 41,097 | 881 (2.14) | 324 (0.79) | 3,557 (8.66) | 36,335 (88.41) | 939 (77.93) | 266 |
| *Cucurbita pepo* | 27,862 | 275 (0.99) | 122 (0.44) | 2,252 (8.08) | 25,213 (90.49) | 269 (67.76) | 128 |
| Total | 194,205 | 2,842 (1.46) | 1,204 (0.62) | 15,908 (8.19) | 174,251 (89.73) | 3,054 (75.48) | 992 |

The R score was broadly distributed across each proteome (median 0.47-0.63), and R alone exceeded 0.5 for 48-60% of proteins (Fig. S1). Integration of domain coherence restricted the High Confidence and Candidate classes to 1.4-2.9% of each proteome (Table S4), showing that the composite classification, rather than the R score alone, provides a selective prioritization.

The Low Confidence/Divergent class (7.65-8.66% across species) comprises proteins with reduced coherence between binary and multi-label outputs, for example, moderate R scores coupled with weak or absent domain signals. These proteins represent the frontier of DREM’s detection capacity and are of secondary interest for immediate functional validation, though they may include genuine resistance genes with highly divergent properties. Detailed results for each species, including absolute counts, percentages for all four confidence classes, and summary statistics are provided in Table S4.

### 2.6. Structural validation via HMM-based domain scanning

To assess the biological validity of DREM predictions and characterize the relationship between sequence-based and domain-based resistance gene annotation, all High Confidence and Candidate proteins identified across the five proteomes were subjected to domain scanning using HMMER v3.3 against Pfam profiles of the five structural domains targeted by the multi-label classifier: NB-ARC (PF00931), TIR (PF01582, PF13676), Kinase (PF00069), LRR (PF00560, PF01462, PF01463), and CC (PF06203, PF03479), as described in Methods. This analysis quantifies the complementarity between sequence-level predictions learned by DREM and traditional domain-based annotation pipelines.

Across all five species, the large majority of High Confidence and Candidate proteins received at least one significant HMMER hit (E-value < 1 × 10̅⁵), confirming the presence of detectable canonical resistance-associated domain architecture. HMMER confirmation rates, the proportion of DREM-predicted candidates with at least one significant HMMER domain hit, ranged from 67.7% in *C. pepo* (269 out of 397 candidates) to 78.0% in *O. sativa* (585 out of 750 candidates), with intermediate values in *V. vinifera* (939/1,205; 77.9%) and *A. thaliana* (811/1,082; 75%) (Table 1). These confirmation rates show substantial concordance between

DREM’s sequence-level signal and structural domain evidence across taxonomically diverse plant species, although DREM does not use domain profiles at inference. The subset of candidate proteins not confirmed by HMMER, designated HMMER-divergent, comprised between 128 (*C. pepo*) and 271 (*A. thaliana*) sequences per species, collectively representing 992 proteins across all five proteomes (Table 1).

These proteins carry a strong DREM resistance signal (High Confidence or Candidate classification) in the absence of detectable canonical domain architecture according to Pfam profiles. Comprehensive classification results including R-score statistics, NLR subclass distributions (TNL, CNL, NLR, RLK/RLP), and HMMER validation metrics for all species are provided in Tables S5 and S6.

Gold candidates were identified from the HMMER-divergent pool by applying stringent thresholds (PRS > 0.75 and R score > 0.90). To annotate their domains with complementary approaches, all gold candidates were submitted to InterProScan against the full InterPro database, which integrates multiple structural databases and prediction tools. This analysis retained 20 curated candidates with InterPro annotations (Fig. 3; Data Set S5).

**Fig. 3.**
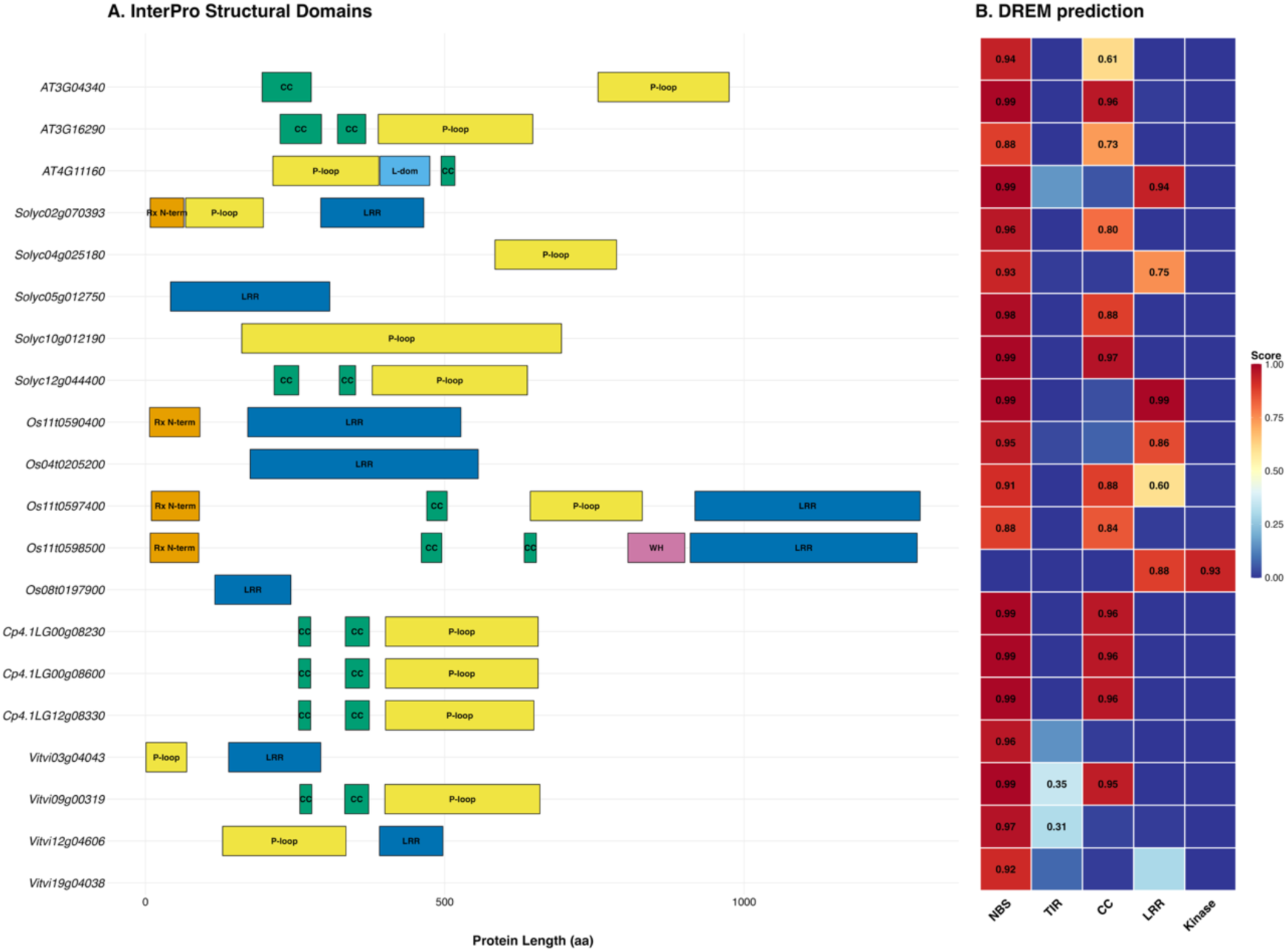
InterProScan validation and DREM domain predictions for 20 curated candidates. **(A)** Structural domain composition detected by InterProScan (multiple integrated tools: Gene3D, SUPERFAMILY, Pfam, SMART and Coils) across the 20 curated candidates, represented as domain architecture cartoons scaled to protein length (aa). Domains are color-coded by type: CC (green), L-dom (cyan), LRR (blue), P-loop (red), Rx N-terminal (brown), WH (purple). **(B)** Corresponding DREM domain probability predictions for the same proteins as a heatmap, with columns representing the five predicted domains (NBS, TIR, CC, LRR, Kinase) and rows representing individual proteins. Color intensity indicates predicted probability (dark blue = high).

In the curated candidates set, 19 of the 20 were assigned by DREM to the CNL or NLR subclasses, and 18 had P(NBS) > 0.88. InterProScan identified domains characteristic of resistance proteins in 8 of the 20 curated candidates: four from *O. sativa* (*Os11t0590400, Os04t0205200, Os11t0597400* and *Os11t0598500*), three from *V. vinifera* (*Vitvi03g04043, Vitvi12g04606* and *Vitvi19g04038*) and one from *S. lycopersicum* (*Solyc02g07039*3), whereas none of the candidates from *A. thaliana* and *C. pepo* carried such domains (Fig. 3; Data Set S5). These annotations included the disease resistance N-terminal domain, found in *Os11t0590400* (567 aa), *Os11t0597400* (1,295 aa), *Os11t0598500* (1,290 aa) and *Solyc02g070393* (466 aa); regions associated with the NB-ARC domain, such as the P-loop, apoptotic protease-activating factor helical and winged-helix domains, in *Solyc02g070393* and *Vitvi03g04043* (293 aa), and an NB-ARC domain-containing annotation in *Vitvi19g04038* (158 aa); and R13L4/SHOC-2-like LRR domains in *Os11t0590400, Os04t0205200* (570 aa) and *Vitvi12g04606* (498 aa). Several of these proteins were short, consistent with truncated or partial resistance-like loci.

The remaining 12 curated candidates carried more general annotations. Nine were P-loop NTPases of the AAA+ ATPase or ABC transporter type, including *AT3G04340*, which combines an AAA+ ATPase with a Peptidase M41 domain, and three paralogous proteins from *C. pepo* (*Cp4.1LG00g08230, Cp4.1LG00g08600* and *Cp4.1LG12g08330*). The others were a GTP-binding protein (AT4G11160), an F-box/LRR protein (*Os08t0197900*) and a protein with an LRR annotation only (*Solyc05g01275*0). These proteins share the P-loop NTPase fold or LRR regions with resistance proteins, which may account for their high DREM scores.

The eight candidates with resistance-associated domains represent priority targets for functional validation, combining strong sequence-level resistance signals with independent domain evidence. Complete DREM predictions and InterProScan annotations for the 20 curated candidates are provided in Data Set S5.

### 2.7. Attention-based interpretability

Attention profiles of the multi-label classifier revealed sharp, localized peaks rather than uniform distributions (Fig. 4), indicating that the model weights specific sequence regions for domain prediction. Enrichment of attention was quantified for nine conserved resistance-associated motifs detected in the 20 curated candidates (Data Set S6). Attention was depleted at the Walker A (P-loop) motif (median enrichment 0.68, IQR 0.49-0.89; n = 10) and the Walker B motif (median 0.17; n = 8), whereas the Kinase-2 motif, detected in all 20 candidates, was enriched in 15 (median 1.71). Attention therefore did not concentrate on the core ATP-binding motifs, consistent with the model drawing on sequence information beyond the conserved signatures captured by domain profiles.

**Fig. 4.**
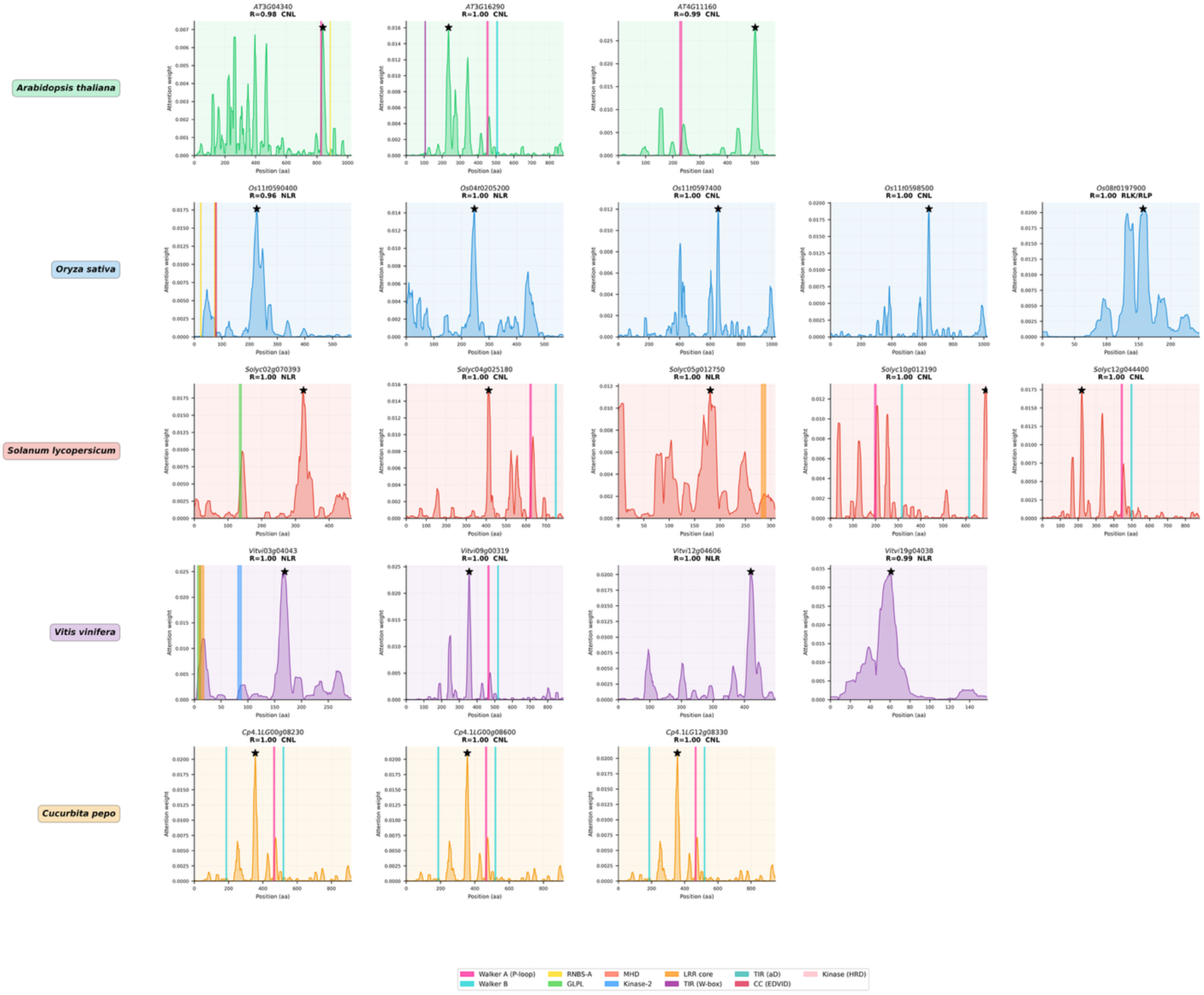
Attention profiles of 20 curated candidates with conserved resistance gene motif annotations. Position-wise attention weights along each protein sequence, shown for the 20 curated candidates and grouped by species (background color). Each panel is labelled with the gene identifier, the DREM resistance score (R) and the predicted structural class (CNL, NLR, RLK/RLP). Colored vertical bands indicate the positions of conserved resistance-associated motifs identified by regular-expression matching (Methods); not all motifs listed in the legend were detected in the curated candidates. The kinase activation-loop motif is shown with the HRD pattern, whereas the enrichment values in Data Set S6 were computed with the DFG pattern, detected only in *AT3G04340*.

To characterize the composition of high-attention regions, amino acid enrichment was calculated within the top 10% of attention-weighted positions of each of the 20 curated candidates (Fig. S2). Leucine was enriched in high-attention regions of all 20 candidates, whereas the aromatic residues phenylalanine and tryptophan were depleted in 19 and 17 of them, respectively, and asparagine and cysteine showed variable patterns depending on protein architecture. Leucine enrichment is consistent with attention to leucine-rich regions, suggesting that, beyond the canonical motifs, the model weights residue composition along the sequence.

### 2.8. Recovery of characterized NLRs in complete proteomes

Experimentally characterized NLRs from RefPlantNLR were matched to their corresponding loci in the proteomes of *A. thaliana*, *S. lycopersicum* and *O. sativa* (33, 10 and 12 loci, respectively; see Methods), and their position in the proteome-wide PRS ranking was examined. These loci were concentrated at the top of each proteome, with median percentile ranks of 1.1% in *A. thaliana*, 1.0% in *S. lycopersicum* and 1.2% in *O. sativa*; 54 of the 55 loci were classified as High Confidence or Candidate and fell within the top 1.7%; the exception was Prf (1,824 aa), which exceeds the 1,024-residue input window (Data Set S7). Since 53 of these loci were absent from the training set, this result reflects the recognition of resistance genes not seen during training. The composite score contributed substantially to this prioritization: ranked by R score alone, the same loci had median percentile ranks of 3.9%, 5.5% and 4.5%, respectively, so that integrating domain coherence placed them on average about four-fold higher in the ranking (Data Set S7).

Because characterized NLRs ranked at the top of each proteome, the proteins ranked alongside them that lack a match among characterized resistance proteins are natural candidates for functional studies. High Confidence proteins of the TNL, CNL and NLR subclasses without a close match in RefPlantNLR or in the PRGdb curated reference collection were therefore ranked by PRS, yielding 133 candidates in *A. thaliana*, 145 in *S. lycopersicum* and 225 in *O. sativa* (Data Set S8). To guide the choice among them, each candidate is reported with its closest characterized resistance protein, the associated pathogen when sequence identity is ≥ 50%, domain support and genomic position. In *S. lycopersicum*, for example, six candidates map to the R2-like cluster on chromosome 4 previously described by Andolfo *et al*. (2014, 2021), all homologous to resistance proteins against *Phytophthora infestans* (Data Set S8).

### 2.9. Comparison with existing resistance-gene predictors

To position DREM relative to existing machine-learning R-gene predictors, its binary classifier was compared with PRGminer (Duhan and Kaundal, 2025), a deep learning predictor that, like DREM, uses protein sequences alone as input, discriminates R-genes from non-R-genes across resistance classes in its first phase, and is available as a standalone tool applicable at proteome scale (Fig. 5A).

**Fig. 5.**
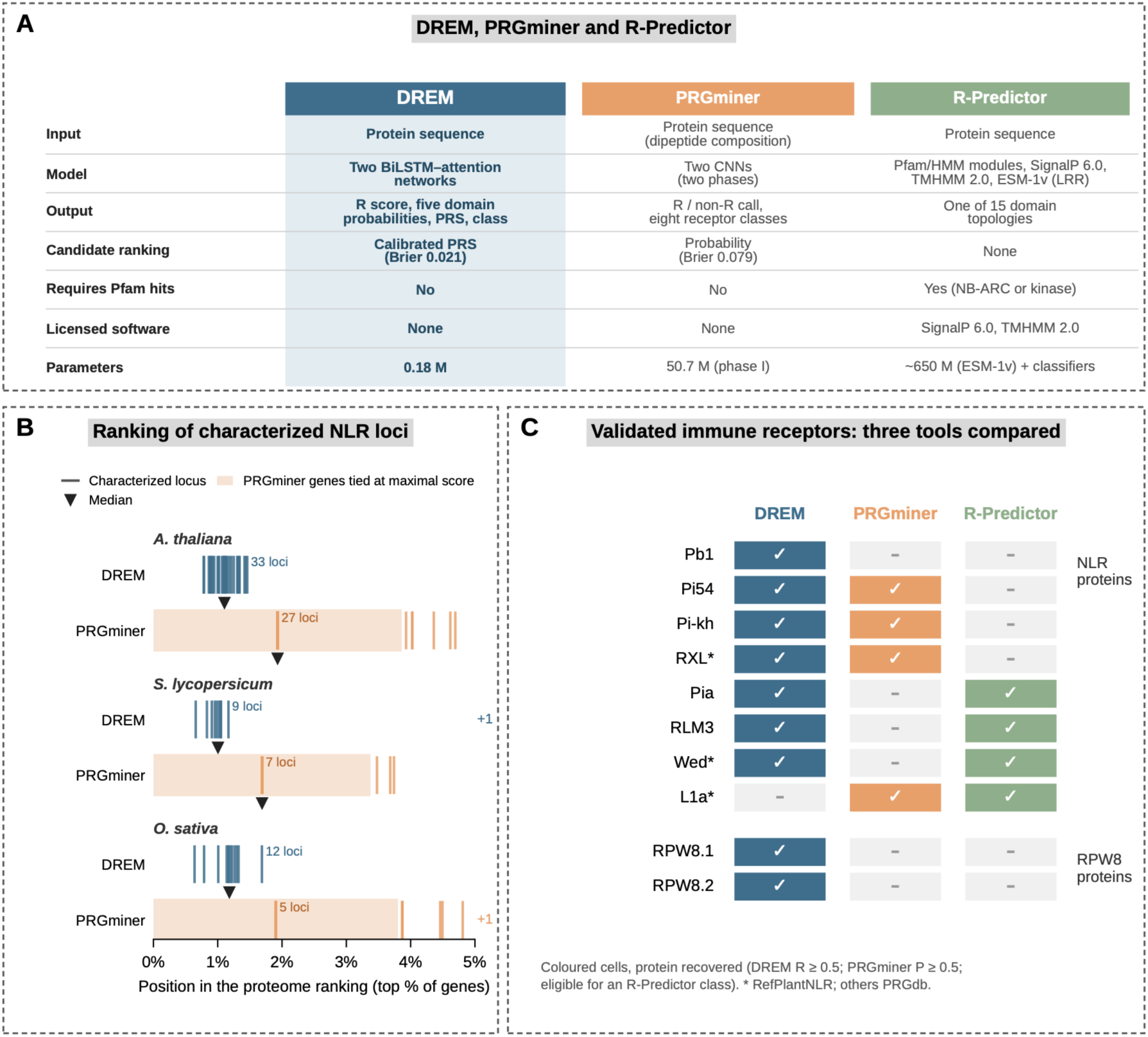
Comparison of DREM with PRGminer and R-Predictor. (A) Main features of DREM, PRGminer and R-Predictor. Brier scores were computed on the held-out test set (n = 15,000). (B) Ranking of the 55 characterized NLR loci in the complete proteomes of *A. thaliana*, *S. lycopersicum* and *O. sativa*. All genes were ranked by DREM (PRS) and by PRGminer (resistance probability); each vertical line marks one locus, positioned as the percentage of the proteome ranked above it, and triangles mark the median. Only the top 5% of each ranking is shown, and loci ranked lower are counted on the right. Shaded areas indicate genes to which PRGminer assigned an identical maximal probability, which cannot be ordered with respect to one another. **(C) Recovery of experimentally validated NLR and RPW8 proteins, excluded from DREM training, for which at least one tool gave a different result.** Colored cells indicate that the protein was recovered: R ≥ 0.5 for DREM, P ≥ 0.5 for PRGminer and, for R-Predictor, eligibility for at least one of its classes based on its Pfam and signal-peptide gating steps (Methods). Asterisks mark RefPlantNLR proteins; the others are from PRGdb.

Both tools were applied to the same 15,000 held-out sequences of the DS150 test set and evaluated at the same decision threshold (0.5) (Table S7; per-protein predictions in Data Set S1). DREM outperformed PRGminer across all metrics: MCC reached 0.938 (95% CI 0.933-0.944) against 0.820 (0.811-0.829), ROC-AUC 0.997 against 0.951, and PR-AUC 0.997 against 0.938. The 95% confidence intervals of the two tools did not overlap for any metric (Table S7), and McNemar’s test confirmed that the two tools differed significantly in their classification errors (1,133 proteins correctly classified only by DREM versus 245 only by PRGminer; p = 3.5 × 10̅¹²⁶). The largest difference concerned false positives: PRGminer assigned 801 non-resistance proteins to the resistance class, 657 of which (82.0%) were correctly rejected by DREM.

Because DREM is intended to rank candidates for experimental validation, the reliability of its continuous scores was also assessed. Among the 1,500 highest-scoring proteins (top 10% of the test set), DREM selected no false positives, compared with 67 for PRGminer; among the top 3,750 (25%), DREM selected one false positive against 166 for PRGminer (Fig. S3B). DREM scores were also better calibrated, with a Brier score of 0.021 against 0.079 for PRGminer (Fig. S3A). PRGminer overestimated resistance probability across intermediate score ranges, whereas DREM tended toward conservative estimates, a desirable property when scores are used to prioritize candidates.

On the complete RefPlantNLR release (481 experimentally characterized NLRs; Kourelis *et al.,* 2021), both tools recovered 478 proteins (99.4%) (Table S7). In complete proteomes, however, PRGminer placed the 55 characterized NLR loci lower than DREM, with median percentile ranks of 1.7-3.9%, because it assigned an identical maximal probability to 1,066-1,362 genes per proteome, which therefore could not be ordered (Fig. 5B; Data Set S7). Among the 20 curated candidates, PRGminer classified 9 as resistance proteins, including all eight candidates with resistance-associated domains, whereas it did not recover any of the candidates from A. thaliana and C. pepo (Data Set S5). As curated candidates were selected from DREM scores, this comparison does not measure DREM accuracy.

R-Predictor (Liu *et al.,* 2026) assigns proteins to its NLR, RLK and TKP modules only when Pfam NB-ARC or kinase domains are detected, and to its RLP and LysM modules only when a signal peptide is predicted; these gating steps were therefore reproduced from its source code (Methods). Because only the gating steps were reproduced, eligibility indicates that a protein could be assigned to an R-Predictor class, not the class it would receive. Among the 148 experimentally validated PRGdb resistance proteins excluded from training, all 119 that could enter an R-Predictor class were also recovered by DREM (R ≥ 0.5), which additionally recovered 19 of the 29 that could not, whereas PRGminer recovered 121 of the 148 proteins. The proteins recovered by DREM but not eligible for R-Predictor included the atypical NLRs *Pb1, Pi-Kh* and *Pi54* and the RPW8 proteins (Fig. 5C), as well as several non-receptor resistance genes, such as *Lr34* and *Xa5*, for which R-Predictor was not designed (Data Set S2). On the 440 RefPlantNLR proteins, which are all NLRs, the two approaches were nearly equivalent: the gating steps excluded only *Pb1* and *RXL*, both recovered by DREM, whereas DREM missed only *L1a* (Data Set S3). Of the 992 HMMER-divergent proteins prioritized by DREM, 945 lacked the features required to enter any R-Predictor class (Data Set S9), as did six of the eight curated candidates with resistance-associated domains (Data Set S5).

Finally, DREM required 183,457 parameters (0.74 MB) against approximately 50.7 million for the PRGminer Phase 1 model (613 MB for both phases) and processed 2,829 sequences per second against 80 on the same GPU (NVIDIA Tesla T4), a 35-fold difference relevant for proteome-scale and database-wide applications (Fig. 5A; Table S7).

## 3. DISCUSSION

The discovery of resistance genes has traditionally relied on homology search tools that identify predefined patterns but lose sensitivity for sequences with divergent regions. DREM is based on learning compositional properties of resistance proteins rather than explicit motif matching. This framework was able to retrieve functionally validated proteins from the RefPlantNLR database (Kourelis *et al.,* 2021) with a sensitivity of 99.8% and an average R score of 0.9934. The model generalized across species, recovering experimentally validated resistance proteins excluded from training and characterized NLR loci in complete proteomes. By integrating resistance and domain probabilities into a composite score, DREM moves from classification to the prioritization of candidates within complete proteomes.

Traditional computational methods, such as those based on HMMs, generally compare a sequence pattern to a target, removing any sequence not strictly conforming to a predefined and rigid profile (Hofberger *et al.,* 2014). In contrast, DREM follows an integrated approach that can capture sequence features distributed along the protein, even when evolutionary pressure has caused significant sequence divergence (Zhang *et al.,* 2024; Hou *et al.,* 2026). Across the five plant proteomes (approximately 194,000 proteins), DREM detected strong resistance signals in 992 proteins lacking canonical domains detectable by HMMER, accounting for 24.5% of High Confidence and Candidate predictions, and 945 of these could not enter any R-Predictor class (Liu *et al.,* 2026). These proteins include candidates with divergent or truncated resistance-associated domains, as well as proteins sharing only general features, such as the P-loop NTPase fold, with resistance proteins, indicating that part of the resistance-protein complement escapes profile-based detection and that complementary annotation remains useful to interpret these candidates.

The technical strength of this framework is rooted in the use of a BiLSTM architecture combined with attention mechanisms that processes amino acid sequences in both directions, capturing long-range dependencies. This capability serves as a foundation, because resistance proteins are modular in nature and their biological function critically depends on long-range interactions between distant domains (Anbu *et al.,* 2023). In contrast, most traditional computational tools analyze sequences through small, local segments or rigid position-weight matrices, limiting their ability to recognize distributed functional signals (Steuernagel *et al.,* 2015; Li *et al.,* 2016). The BiLSTM overcomes this constraint by creating a vector representation of the whole sequence that is less dependent on the exact position of conserved motifs, a property relevant for gene families shaped by duplication and rearrangement (Berman et al., 2026). In line with this, attention analysis showed that the model does not concentrate on the core ATP-binding motifs, and the enrichment of leucine in high-attention regions points to attention to leucine-rich regions, suggesting that DREM draws on sequence information beyond the conserved signatures captured by domain profiles.

This ability to handle diverse sequence contexts is necessary but not sufficient for reliable proteome-wide prediction. Because resistance proteins represent only a small fraction of each proteome, even a low false-positive rate produces many spurious candidates when a model is applied genome-wide (Ding *et al.,* 2024). To address it, DREM integrates two different scores, the R score and the SCS. While the R score evaluates the primary sequence logic, the SCS assesses the coherence of the predicted domain architecture, and the PRS combines the two. This is relevant for highly variable domains such as the coiled-coil domain, whose natural structural diversity often underpins functional variation (Sulkowski *et al.,* 2025; Wang *et al.,* 2026). Although the CC domain shows a lower MCC (0.701) compared to the more conserved NBS (0.952) or Kinase domains, DREM’s scoring system transforms this natural biological diversity into a quantitative confidence parameter. In practice, the composite classification excluded all UniProt negative controls from the High Confidence and Candidate classes, although 99 of them had R ≥ 0.5, and placed characterized NLR loci 3.5-to 5.5-fold higher in proteome rankings than the R score alone. This makes it possible to move beyond binary classifications and prioritize candidates for experimental validation based on multiple lines of evidence.

The comparison with existing predictors supports this design. On identical held-out data, DREM reduced classification errors relative to PRGminer (Duhan and Kaundal, 2025) by approximately two-thirds, mainly by reducing false positives, which represent the main cost of proteome-wide screening because each spurious candidate translates into unnecessary validation effort. Its scores were also better calibrated, so that the R score and the derived PRS can be used to prioritize candidates rather than only to label them. As the held-out set shares the data distribution used to train DREM, the comparisons on independent data are more informative. On the canonical NLRs of RefPlantNLR (Kourelis *et al.,* 2021), DREM, PRGminer and the R-Predictor gating steps performed similarly, suggesting that the advantage of DREM lies less in recognizing well-characterized architectures than in separating and ranking resistance proteins within the rest of the proteome. In complete proteomes, 54 of the 55 characterized NLRs ranked within the top 2% of genes by PRS, whereas PRGminer assigned an identical maximal probability to more than a thousand genes per proteome, which could not be ordered. Beyond canonical receptors, DREM also recovered experimentally validated proteins that cannot enter any R-Predictor class (Liu *et al*., 2026), including the atypical NLRs Pb1, Pi-Kh and Pi54, RPW8 proteins and non-receptor resistance genes such as Lr34 and Xa5. Finally, the compact architecture of DREM, with about 277-fold fewer parameters than PRGminer, facilitates its integration into resources such as PRGdb (Calle García *et al.,* 2022), where large numbers of sequences must be scored and periodically re-annotated.

Among the candidates that escape profile-based detection, the eight curated candidates with resistance-associated domains, from rice, grapevine and tomato, represent the most immediate targets for functional studies. Several of them are short proteins, and they may represent atypical receptors or truncated variants located in genomic regions where frequent rearrangements make standard annotation difficult (Sutherland *et al.,* 2024; Belinchon-Moreno *et al.,* 2025). Ranking complete proteomes by PRS also highlighted high-confidence NLRs without characterized counterparts. In tomato, six of these candidates map to the R2-like cluster on chromosome 4, the largest NLR cluster of the tomato genome, which has been proposed to harbor functional resistance genes not yet identified (Andolfo *et al.,* 2014, 2021). These candidates are homologous to resistance genes against Phytophthora infestans such as R2 and Rpi-blb3. This convergence between an independent genomic analysis and DREM prioritization indicates that the ranking can direct functional studies towards regions already known to be involved in disease resistance.

These results suggest that part of the natural diversity of immune receptors available for crop improvement escapes homology-based annotation (Förderer and Kourelis, 2023), and that combining profile-based and sequence-based approaches can help to recover it for breeding and sustainable agriculture. In conclusion, DREM extends R-gene discovery beyond homology by integrating resistance and domain probabilities learned from sequence. It does not aim to replace established methods but serves as a complementary tool for genomic annotation in the big data era (Song *et al.,* 2025; Dublino and Ercolano, 2025): lightweight and applicable to complete proteomes, it provides a calibrated ranking that links genome-wide prediction to the selection of candidates for functional validation, including atypical receptors missed by domain-based annotation. Such tools are becoming increasingly important as pathogens evolve rapidly under the pressure of global environmental changes (Ercolano *et al.,* 2022; Singh *et al.,* 2023; Kumar and Mukhopadhyay, 2025). Integrating deep learning into plant genomics can therefore help to exploit the full potential of natural immunity for the development of resilient crops.

## 4. METHODS

### 4.1. Dataset and preprocessing

The training dataset was constructed from two complementary sources. Putative resistance proteins were extracted from PRGdb 4.0 (Calle García *et al.,* 2022), a manually curated reference database for plant resistance genes developed and maintained by our research group. Non-resistance sequences were retrieved from UniProt (Ahmad *et al.,* 2025; The UniProt Consortium, 2025), selecting protein sequences lacking any specific annotation associated with resistance mechanisms. For each protein, the amino acid sequence and two types of binary labels were recorded: a general resistance label (1 for resistant, 0 for non-resistant), and five domain-specific labels (NBS, LRR, TIR, CC, kinase) indicating the presence or absence of principal structural domains associated with plant resistance genes. The initial dataset comprised 586,652 putative resistance protein sequences and an equivalent number of non-resistance sequences from UniProt, totaling approximately 1.17 million protein sequences. Before training, the dataset went through multi-step quality control: sequences containing non-standard amino acid characters were removed, incomplete records and sequences with missing or ambiguous annotations were excluded, and exact duplicates were removed, with priority assigned to resistance-annotated entries whenever identical sequences carried conflicting labels. A second deduplication step was applied after the train/validation/test split to prevent sequence leakage across subsets, without applying sequence similarity clustering thresholds. The final curated dataset comprised 956,576 unique protein sequences and was used as the source population for model development, from which balanced random subsets of 50,000, 100,000 and 150,000 sequences (DS50, DS100 and DS150) were sampled to evaluate the effect of training-set size on convergence, predictive performance, and generalization.

### 4.2. Sequence encoding and model architecture

Protein sequences were encoded as integer sequences, assigning each amino acid a unique index (1-20) and reserving index 0 for padding. Sequences longer than 1,024 residues were truncated to this length. Since 99% of sequences in the dataset were shorter than 891 residues, truncating at 1,024 preserved nearly all biological data while limiting computational cost. Shorter sequences were padded to match the batch maximum using a custom function that prevented the model from computing gradients on padding positions.

Integer-encoded sequences were passed through an embedding layer that converts each amino acid index into a vector of 32 numbers. Embedding sizes of 16, 32, and 64 were compared on the validation set by ROC-AUC, and 32 gave the best performance. The embedding layer was initialized with random weights and trained alongside the rest of the network.

DREM consists of two networks with identical architecture, trained separately. The first is a binary classifier predicting whether a sequence belongs to the resistance class (R score), representing the model’s confidence in resistance function. The second is a multilabel classifier predicting the presence or absence of five structural domains: NBS, LRR, TIR, CC and kinase. The two networks differ only in the dimensionality of the output layer.

The core architecture is a bidirectional Long Short-Term Memory network (BiLSTM) following Hochreiter and Schmidhuber (1997) for LSTM and Schuster and Paliwal (1997) for the bidirectional extension, implemented in PyTorch (v2.0.1, Paszke *et al.,* 2019) with additive attention pooling. The BiLSTM processes sequences across two stacked layers, each with 64 units per direction, producing a 128-dimensional hidden representation at each position Eq. (1a-1c):

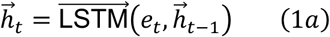

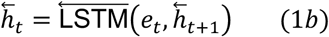

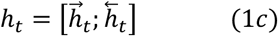

where *e_t_*is the embedded token (amino acid vector) at position *t*, ℎ^#⃗^*_t_*is the hidden state from the forward LSTM, ℎ⃖^#^*_t_*is the hidden state from the reverse LSTM, and ℎ*_t_*denotes concatenation of the two directions. Position-wise hidden states are aggregated through an additive attention pooling layer following Bahdanau *et al.,* 2014, Eq. (2a-2b):

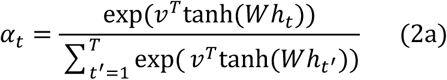

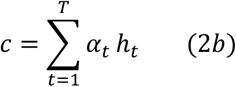

where *w*is a learnable weight matrix that transforms the hidden state, *v*is a learnable context vector that scores positional importance, *α_t_*is the normalized attention weight for position *t*, ℎ*_t_*is the hidden state at position *t*, *T* is the sequence length and *c* is the final 128-dimensional context vector summarizing the sequence. The attention scores sum to 1, and softmax is applied only to non-padding positions. This summary vector is fed to a linear layer (128 units), followed by ReLU activation, dropout (p = 0.1) and a final output layer. For the binary classifier, the resistance probability *R* is computed as defined in Eq. (3):

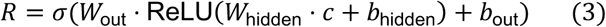

where *c* is the context vector, *w*_hidden_and *b*_hidden_are the weights and bias of the first linear layer (128 units), *w*_out_and *b*_out_are the weights and bias of the final output layer, and *σ*is the sigmoid function that converts the raw output to a probability. For the multi-label classifier, five independent outputs are produced, and each is converted to a probability. Each network has approximately 184,000 trainable parameters.

### 4.3. Training procedure

Training configurations were generated by balanced random sampling from the full source population. Different dataset sizes (50,000, 100,000, and 150,000 sequences) and multiple epoch durations (10, 20, 30, and 50 epochs) were tested. The 150,000-sequence setting trained for 10 epochs emerged as optimal based on validation loss convergence patterns and downstream classification performance (Table S1), representing a good trade-off between overfitting and underfitting given the dataset size. To ensure robust assessment of model generalization, 5-fold stratified cross-validation was applied on the 150,000-sequence dataset. For each fold, the data was partitioned into training (72%, n=108,000), validation (8%, n=12,000), and test (20%, n=30,000) sets, with stratification applied to maintain class balance. A fixed random seed (42) was used throughout all random operations to ensure reproducibility across different hardware platforms and library versions. Performance metrics (accuracy, F1-score, AUC) were computed on each fold’s independent test set, with final reported values representing the mean ± standard deviation across all five folds (Table S2).

Both classifiers were trained using the Adam optimizer (Kingma and Ba, 2014) with learning rate 1×10̅³ and binary cross-entropy loss applied independently to each output unit in the multi-label setting, as defined in Eq. (4):

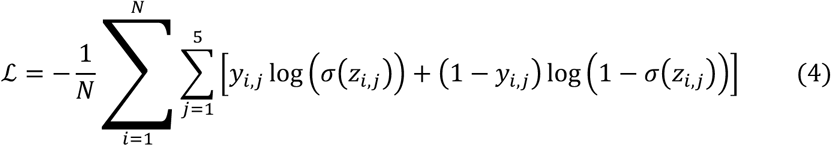

where *N* is the batch size, *y_i_*_,j_ is the ground truth label for domain j in sequence *i*, *z_i_*_,j_ is the raw logit output, and *σ* is the sigmoid function.

Each domain is treated independently as a binary problem, since domains can occur in different combinations across resistance genes. Gradient clipping (maximum norm 1.0) ensured stable training and prevented gradient explosions. Batch size was set to 32 with dynamic padding at the batch level. Model checkpointing saved the state with lowest validation loss, which was then evaluated on the held-out test set of 15,000 sequences (10% of DS150). All experiments were conducted on an NVIDIA Tesla T4 GPU using Python v3.10, NumPy v1.24.3 (Harris *et al.,* 2020), and scikit-learn (v1.3.0, Pedregosa *et al.,* 2011).

### 4.4. Evaluation metrics and external validation

Model reliability was assessed through internal evaluation on the held-out test set, and external validation on independent datasets. For the binary classifier, accuracy, ROC-AUC, Matthews Correlation Coefficient (MCC), F1-score, precision and recall were computed at a decision threshold of 0.5. For each domain, ROC-AUC, Average Precision (PR-AUC), MCC, F1-score, precision, and recall were computed. At the multi-label level, Exact Match Accuracy and Hamming Loss were computed. Bootstrap resampling (1,000 iterations) on the test set provided 95% confidence intervals for ROC-AUC and MCC.

Two independent validation analyses were performed on datasets not used during model development to ensure unbiased assessment of generalization. The first validation set comprised 152 resistance proteins from PRGdb 4.0 with experimentally validated resistance function reported in primary literature, of which 148 were retained after sequence-level deduplication, balanced with 152 non-resistance proteins from UniProt. The second validation set consisted of 440 sequences from RefPlantNLR (Kourelis *et al.,* 2021), a manually curated database of experimentally characterized plant NLR proteins, balanced with 440 non-resistance proteins from UniProt. In both cases, resistance proteins from external databases and non-resistance proteins from UniProt were deliberately excluded from the training dataset to ensure unbiased validation and prevent data leakage. Both validation sets were processed through the standard inference pipeline. Sensitivity was computed as the proportion of resistance proteins correctly predicted (above the 0.5 threshold), and specificity as the proportion of non-resistance proteins correctly predicted (below the 0.5 threshold). Recovery at R ≥ 0.8 was additionally reported as a high-confidence threshold (Data Sets S2 and S3).

### 4.5. Inference scoring framework

Each protein sequence passing through the model produces two outputs: a Resistance Score (R) from the binary classifier and domain probabilities from the multi-label classifier. These raw outputs are integrated into biologically interpretable confidence scores. The R score represents the model’s confidence that the sequence belongs to the resistance gene class, ranging from 0 to 1, based on its similarity to training data.

The Structural Confidence Score (SCS) evaluates whether predicted domains form the typical structural patterns of functional resistance proteins and is calculated according to Eq. 5:

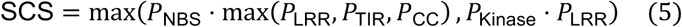

where *P*_NBS_, *P*_LRR_, *P*_TIR_, *P*_CC_, *P*_Kinase_are the predicted probabilities (0-1) for each domain from the multi-label classifier, and maxselects the highest value. For NLR-type proteins, the first term (*P*_NBS_ ⋅ max(*P*_LRR_, *P*_TIR_, *P*_CC_)) scores NBS paired with whichever accessory domain (LRR, TIR, or CC) is most strongly predicted for RLK-type proteins, the second term (*P*_Kinase_ ⋅ *P*_LRR_) scores Kinase paired with LRR. Two additional terms score RLP-type profiles (R · *P*_LRR_, when R > 0.90, *P*_LRR_> 0.80, *P*_NBS_ < 0.30 and *P*_Kinase_< 0.30) and kinase-only profiles (R · *P*_Kinase_, when R > 0.90, *P*_Kinase_> 0.85, *P*_LRR_< 0.30 and *P*_NBS_ < 0.30). The final SCS takes the maximum of these terms.

Both R and SCS provide complementary biological information, so they are combined into a single composite metric. The Protein Resistance Score (PRS) integrates both signals through Eq. 6:

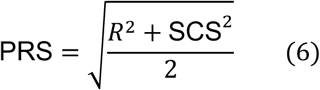

High PRS indicates strong agreement between both signals. However, both signals provide independent biological information. A sequence with high R but lower SCS represents a divergent R-gene with atypical domain composition, where the sequence-level features still suggest resistance. A sequence with high SCS but lower R represents a sequence with perfect domain coherence but different sequence patterns, where structural coherence confirms plausibility. Proteins were stratified into four confidence classes based on different thresholds. Proteins were assigned to confidence classes by hierarchical rules combining R and the domain probabilities. Proteins with a canonical domain profile, namely NLR (*P*_NBS_ > 0.60 and R > 0.60), RLK (*P*_Kinase_ > 0.85, *P*_LRR_ > 0.40 and R > 0.80), RLP or kinase-only (as defined above), were classified as High Confidence when PRS > 0.75 and as Candidate otherwise; NLR proteins were further assigned to the TNL (*P*_TIR_ > 0.45), CNL (*P*_CC_> 0.40) or NLR subclass. Proteins with R > 0.90 lacking canonical domain support, or with R > 0.40 and partial domain signals, were classified as Low Confidence/Divergent. All remaining proteins were classified as Non-resistance-like.

### 4.6. Proteome-wide inference

Proteome-wide inference was performed on complete proteomes of five plant species that were retrieved from Ensembl Plants (Yates *et al.,* 2022) with the following assembly versions: *Arabidopsis thaliana* (TAIR10 assembly, Araport 11 annotation, Cheng *et al.,* 2017), *Oryza sativa* (IRGSP-1.0, Kawahara *et al.,* 2013), *Solanum lycopersicum* (SL3.0, The Tomato Genome Consortium, 2012), *Vitis vinifera* (PN40024.v4, Velt *et al.,* 2023), and *Cucurbita pepo* (v4.1, Montero-Pau *et al.,* 2018).

Proteomes were processed without pre-filtering except for removal of sequences with non-standard amino acids, maintaining consistency with training data preparation.

Each protein sequence was passed through the two-step classification pipeline. The R score and domain probabilities were integrated using the scoring framework described above to calculate SCS and PRS for each protein. The inference was performed without species-specific adaptation in order to verify the model’s ability to generalize to unseen proteomes.

High-confidence candidates were identified using PRS > 0.75, representing proteins with consistent signals across both classifiers. Proteins with PRS > 0.75 and R > 0.90 were designated as gold candidates, representing the most reliable predictions where both the global resistance signal and structural domain coherence align with high confidence. These gold candidates were selected for downstream structural validation and interpretability analysis.

### 4.7. Structural validation and attention analysis

Structural validation was performed on all predicted candidates across the five plant proteomes using domain-based scanning. HMMER v3.3 (Eddy, 2011; Potter *et al.,* 2018) was used to search against Pfam release 36.0 (Paysan-Lafosse *et al.,* 2025) for five target domains: NB-ARC (PF00931), TIR (PF01582, PF13676), Kinase (PF00069), LRR (PF00560, PF01462, PF01463), and CC (PF06203, PF03479). Pfam does not provide a dedicated profile for plant NLR coiled-coil domains, so HMMER confirmation represents a conservative estimate. Candidates were classified as HMMER-confirmed (at least one significant domain hit, E-value < 1 × 10̅⁵) or divergent (no canonical domain detected). This partitioning quantified the complementarity between sequence-level predictions and traditional domain-based annotation.

Gold candidates (PRS > 0.75 and R > 0.90) within the divergent pool were further annotated using InterProScan (v5.59, Jones *et al.,* 2014) against InterPro (release 95.0, Blum *et al.,* 2025) for comprehensive domain annotation. Proteins with at least one InterPro entry providing structural or functional evidence consistent with resistance-related function, including NTPase, coiled-coil, leucine-rich repeat, and kinase domain annotations, were designated as curated candidates and retained for downstream analysis.

To examine which sequence regions contribute to the predictions, position-wise attention weights were extracted for the 20 curated candidates, mapped onto the original sequence and visualized as position-wise heatmaps using matplotlib (v3.7.1, Hunter, 2007) and seaborn (v0.12.2, Waskom, 2021). Attention enrichment ratios were computed for conserved sequence motifs identified by regular expression matching using established consensus patterns from NBS-LRR resistance genes. Motif positions shown in Fig. 4 were obtained with eleven consensus patterns (Walker A, Walker B, RNBS-A, GLPL, MHD, Kinase-2, LRR core, TIR W-box, TIR αD, CC EDVID and Kinase HRD). The enrichment analysis included the nine motifs detected among the 20 curated candidates (Walker A, Walker B, RNBS-A, GLPL, Kinase-2, LRR core, TIR W-box, CC EDVID and Kinase DFG), in which the kinase activation-loop motif was matched with the DFG pattern (Data Set S6). Motif-level enrichment was calculated according to Eq. (7):

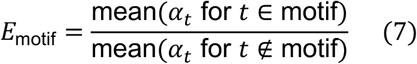

where *α_t_*is the attention weight at position *t*. Values above 1.0 indicate preferential attention to the specific region; values below 1.0 indicate depletion (lower attention). Amino-acid enrichment was calculated for each protein as the frequency of each residue within the top 10% of attention-weighted positions relative to its frequency in the remaining positions (Fig. S2).

### 4.8. Recovery of characterized NLRs and candidate selection

Characterized NLR loci were identified by aligning the 440 independent RefPlantNLR proteins to the five proteomes with DIAMOND (v2, --more-sensitive, Buchfink *et al.,* 2021), retaining the best hit with ≥ 90% identity and ≥ 80% coverage of the RefPlantNLR protein. Matching loci were found only in *A. thaliana, S. lycopersicum* and *O. sativa*. Proteome-wide rankings were computed at the gene level, with each gene represented by its highest-scoring isoform, and loci whose sequences were identical to sequences of the DS150 training split were flagged. For each gene, the percentile rank was calculated as its position in the proteome-wide ranking divided by the total number of genes, separately for PRS and R score.

Candidate NLRs were selected among High Confidence proteins of the TNL, CNL and NLR subclasses. Proteins with ≥ 90% identity and ≥ 80% coverage to any protein of RefPlantNLR or of the PRGdb curated reference collection were considered already characterized and excluded. For the remaining candidates, the best hit (E-value ≤ 10̅⁵) in the PRGdb curated collection was reported as the closest characterized homolog, together with its associated pathogen when identity was ≥ 50%. Genomic coordinates were taken from Ensembl Plants annotations, and the number of other genes classified as High Confidence or Candidate within 200 kb was recorded.

### 4.9. Comparison with existing tools

Comparison tools were selected according to four criteria: peer-reviewed publication, sequence-only input, applicability to the binary discrimination of resistance versus non-resistance proteins across R-gene classes, and availability of a standalone implementation suitable for proteome-scale application. PRGminer met all criteria and was compared with DREM on its predictions. R-Predictor, which assigns proteins to discrete domain topologies without a resistance score, was compared through its gating steps.

#### 4.9.1. PRGminer

PRGminer v0.1.0 (Duhan and Kaundal, 2025) was obtained from its public repository (https://github.com/usubioinfo/PRGminer) and run in Phase 1 mode (R-gene versus non-R-gene) with default parameters. For compatibility with the current TensorFlow release, model loading was redirected to tf-keras (v2.21) without any modification of model architecture or weights. PRGminer probabilities, reported as percentages, were rescaled to [0,1], and both tools were evaluated at a decision threshold of 0.5 on the DS150 held-out test set (15,000 sequences: 7,542 resistance and 7,458 non-resistance proteins) and on the complete RefPlantNLR release (481 sequences). The complete release was used here, rather than the 440-sequence set described above, so that neither tool benefited from dataset filtering. PRGminer processed sequences in batches of 1,000, as recommended by its authors. For DREM inference on RefPlantNLR, sequences were processed individually (batch size 1) and truncated at 1,024 residues, as in training; 262 of the 481 sequences exceeded this length. Accuracy, precision, recall, F1-score, MCC, ROC-AUC, PR-AUC and Brier score were computed with scikit-learn, and calibration was assessed with reliability diagrams over ten equal-width probability bins. To evaluate ranking, proteins were sorted by predicted probability, and the cumulative number of false positives was recorded at each rank, with ties resolved at random over 100 repetitions. 95% confidence intervals for each metric were estimated from 1,000 paired bootstrap resamples of the test set, and differences in classification errors were assessed with McNemar’s test with continuity correction (McNemar, 1947). Model size was measured as the number of trainable parameters and checkpoint file size; throughput was measured end-to-end, including model loading, on 2,000 test sequences using the same NVIDIA Tesla T4 GPU. PRGminer was also applied, with the same settings, to the 148 PRGdb and 440 RefPlantNLR validation proteins and to the complete proteomes of A*. thaliana, S. lycopersicum* and *O. sativa*, after restricting sequences to the 20 standard amino acids. Proteome-wide rankings were computed at the gene level as for DREM, and genes with identical PRGminer probability received the average rank of the tied block.

#### 4.9.2. R-Predictor

R-Predictor (Liu *et al.,* 2026) assigns proteins to its NLR, RLK and TKP modules only when Pfam NB-ARC or protein kinase domains are detected, and to its RLP and LysM modules only when a signal peptide is predicted; downstream modules (ESM-LRR, TMHMM, TIR/RPW8 and coiled-coil detection) only subdivide proteins that pass these gates. The gating steps were therefore reproduced from its source code (https://github.com/zhouyflab/R-Predictor, commit 78a49eb, accessed October 2026). Proteins were scanned with hmmscan (HMMER v3.3.2) against the Pfam profiles distributed with R-Predictor (Pkinase PF00069, PK_Tyr_Ser-Thr PF07714, NB-ARC PF00931, TIR PF01582, PF13676 and PF18567, RPW8 PF05659 and LysM PF01476) using gathering thresholds, as in the pfam_scan.pl call of the original pipeline. Because the R-Predictor profile database contains only these families, hits to the kinase (CL0016) and P-loop NTPase (CL0023) clans used by the original scripts correspond to the Pkinase/PK_Tyr_Ser-Thr and NB-ARC profiles. Twenty-eight of the 992 HMMER-divergent proteins had NB-ARC hits with the R-Predictor profiles and gathering thresholds, reflecting differences in profile version and significance threshold relative to the HMMER scan described above. Proteins lacking NB-ARC and kinase hits were analyzed with SignalP 6.0 (Teufel et al., 2022; eukarya, fast mode), as in R-Predictor. Proteins lacking NB-ARC and kinase hits and a predicted signal peptide cannot be assigned to any R-Predictor class; the final class of proteins passing the gates was not determined. The gating steps were applied to the 992 HMMER-divergent proteins, the 20 curated candidates, the 148 PRGdb validation proteins and the 440 RefPlantNLR proteins (Data Sets S2, S3, S5 and S9). PRGdb proteins recovered by DREM but not eligible for R-Predictor were grouped as atypical NLRs, RPW8 proteins or non-receptor resistance genes according to their literature annotation in PRGdb.

## DATA AVAILABILITY

Resistance proteins used for model development were obtained from PRGdb 4.0 (https://prgdb.org/) and non-resistance proteins from UniProt (https://www.uniprot.org/). External validation sets were derived from experimentally validated PRGdb 4.0 entries and RefPlantNLR (Kourelis et al., 2021). Complete plant proteomes used for proteome-wide inference were obtained from Ensembl Plants (https://plants.ensembl.org/) with the following assembly versions:

- *Arabidopsis thaliana*: TAIR10 (https://plants.ensembl.org/Arabidopsis_thaliana)
- *Oryza sativa*: IRGSP-1.0 (https://plants.ensembl.org/Oryza_sativa)
- *Solanum lycopersicum*: SL3.0 (https://plants.ensembl.org/Solanum_lycopersicum)
- *Vitis vinifera*: PN40024.v4 (https://plants.ensembl.org/Vitis_vinifera)
- *Cucurbita pepo*: v4.1 (https://plants.ensembl.org/Cucurbita_pepo)

PRGminer and R-Predictor were obtained from their public repositories (Methods). Per-protein DREM predictions for the held-out test set, the validation sets, the proteome-wide analyses and the comparisons with PRGminer and R-Predictor are provided in Supplementary Data Sets S1–S9. Dataset composition and preprocessing are described in Methods.

## CODE AVAILABILITY

DREM inference code and trained models are deposited at https://github.com/Rita-Dublino/DREM_ and will be made publicly available upon publication in a peer-reviewed journal; they are available upon reasonable request.

## AUTHOR CONTRIBUTIONS

**R.D.:** Data curation, Methodology, Investigation, Formal analysis, Validation, Writing – original draft. **V.C.:** Methodology, Formal analysis. **G.S.:** Supervision, Writing – review and editing. **M.E.:** Conceptualization, Supervision, Writing – review and editing, Funding acquisition.

