## Supplemental Information for "DREM: integrating resistance and domain probabilities to prioritize plant immune receptors"

### Supplementary Figures

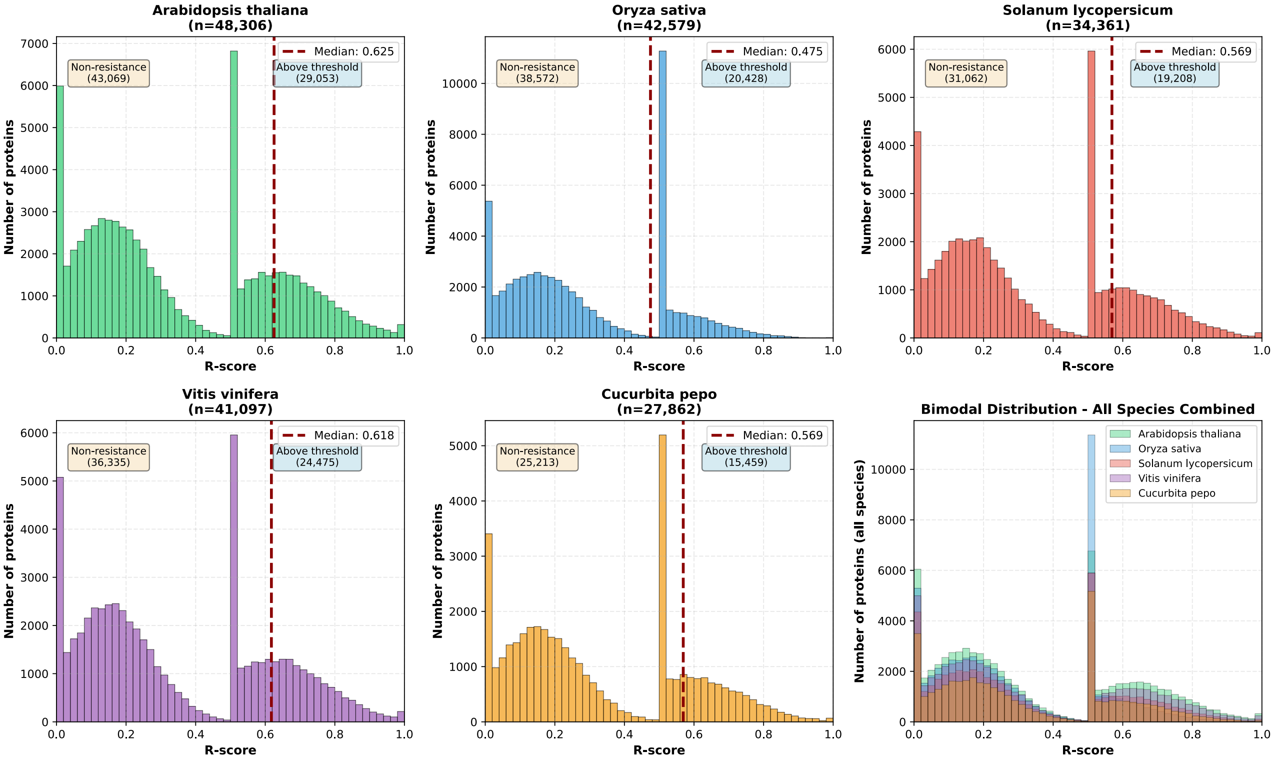

**Supplementary Figure S1. R-score distributions across five plant proteomes.** Each panel shows the R-score histogram for one species; the bottom-right panel overlays all species. Red dashed lines indicate median R scores. Counts above threshold refer to proteins with R > 0.5, before stratification into confidence classes.

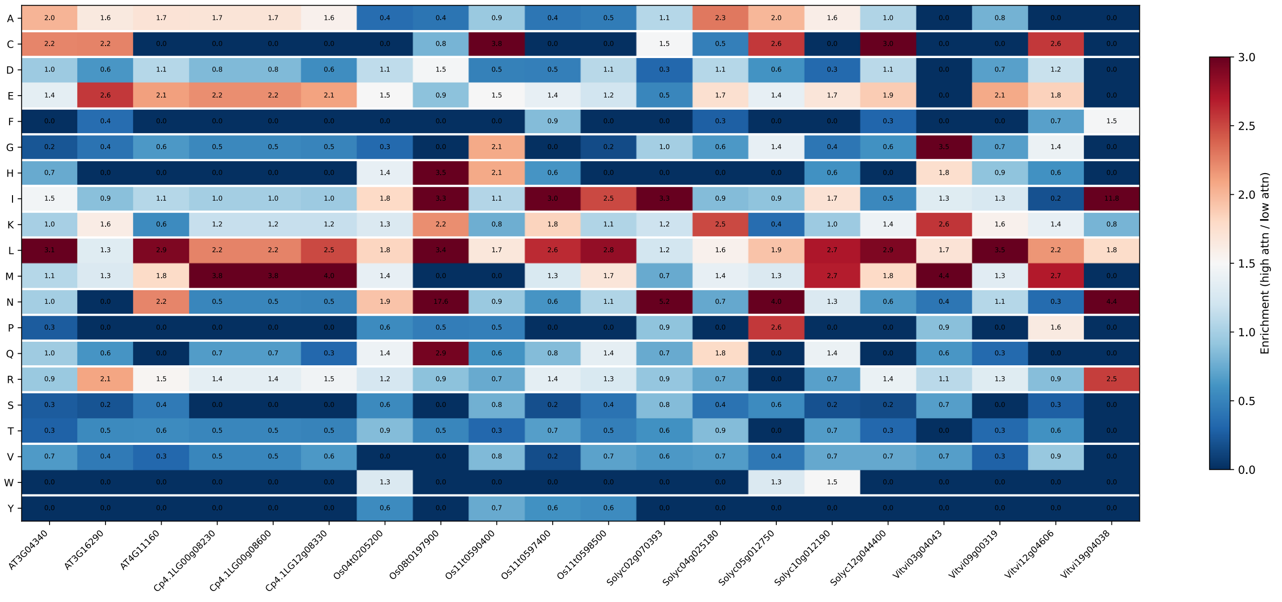

**Supplementary Figure S2. Amino-acid enrichment in high-attention regions of the curated candidates.** Heatmap of the enrichment ratio (red = enriched, blue = depleted, relative to 1.0) of each of the 20 standard amino acids in the top 10% of attention-weighted positions for each of the 20 curated candidates, grouped by species.

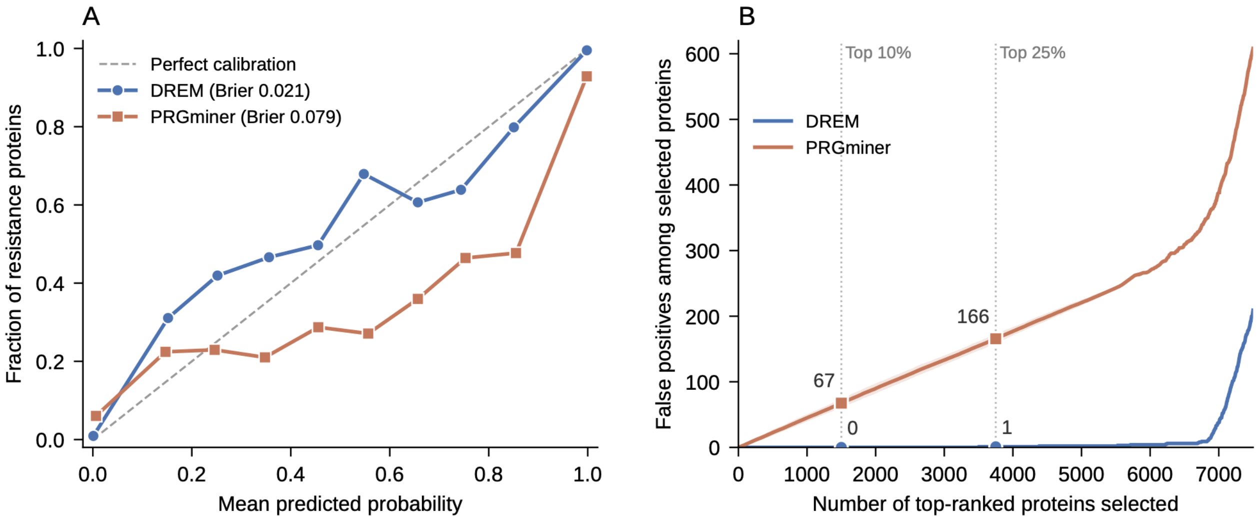

**Supplementary Figure S3. Calibration and ranking of DREM and PRGminer scores on the held-out test set, related to Fig. 5A.**

**(A)** Reliability diagram showing the fraction of resistance proteins in ten equal-width bins of predicted probability; the dashed line indicates perfect calibration.

**(B)** Cumulative number of false positives among the top-ranked proteins (n = 15,000). Lines show the mean and shaded bands ±1 SD over 100 random orderings of tied scores; PRGminer assigns its maximal score to 5,568 proteins and DREM to 74. Dotted lines mark the top 10% (1,500 proteins) and top 25% (3,750 proteins) of the ranking.

#### Supplementary Tables

**Supplementary Table S1. Performance of the binary and multilabel classifiers across training dataset sizes.**

For the binary classifier: accuracy, ROC-AUC, MCC, F1, precision, and recall computed at a decision threshold of 0.5. For the multi-label classifier: Exact Match, Hamming Loss, Micro F1, Macro F1, macro-averaged ROC-AUC, macro-averaged PR-AUC, and macro-averaged MCC. Balanced sampling was applied in all configurations (1:1 ratio of resistant and non-resistant proteins). The model checkpoint corresponding to the lowest validation loss was used for test set evaluation in all conditions. DS150 (bold) was selected as the final configuration.

| | **Binary classifier** | | | | | |
|---|---|---|---|---|---|---|
| **Dataset** | **Accuracy** | **ROC-AUC** | **MCC** | **F1** | **Precision** | **Recall** |
| DS50 | 0.9610 | 0.9896 | 0.9222 | 0.9609 | 0.9708 | 0.9512 |
| DS100 | 0.9646 | 0.9953 | 0.9294 | 0.9648 | 0.9545 | 0.9753 |
| **DS150** | **0.9691** | **0.9967** | **0.9383** | **0.9691** | **0.9743** | **0.9641** |

| | **Multi-label classifier** | | | | | | |
|---|---|---|---|---|---|---|---|
| **Dataset** | **Exact Match** | **Hamming Loss** | **Micro F1** | **Macro F1** | **Macro ROC-AUC** | **Macro PR-AUC** | **Macro MCC** |
| DS50 | 0.9162 | 0.0183 | 0.93 | 0.86 | 0.987 | 0.907 | 0.854 |
| DS100 | 0.9279 | 0.0156 | 0.94 | 0.90 | 0.992 | 0.941 | 0.888 |
| **DS150** | **0.9367** | **0.0136** | **0.95** | **0.90** | **0.992** | **0.940** | **0.893** |

**Supplementary Table S2. DREM classifier performance in 5-fold stratified cross-validation.**

Results from 5-fold stratified cross-validation on 150,000 balanced sequences (75,000 resistance, 75,000 non-resistance), showing fold-wise and mean metrics (accuracy, F1-score, AUC) for both the binary resistance classifier and the multi-label domain classifier (predicting NBS, LRR, TIR, CC, Kinase domains). Each fold used 108,000 sequences for training, 12,000 for validation, and 30,000 for testing. Standard deviations indicate model robustness across independent data partitions.

| | **Binary model** | | | **Multilabel model** | | |
|---|---|---|---|---|---|---|
| **Fold** | **Accuracy** | **F1-Score** | **AUC** | **Accuracy** | **F1-Score** | **AUC** |
| **1** | 0.9731 | 0.9731 | 0.9970 | 0.9374 | 0.9455 | 0.9967 |
| **2** | 0.9699 | 0.9701 | 0.9969 | 0.9398 | 0.9447 | 0.9965 |
| **3** | 0.9707 | 0.9708 | 0.9973 | 0.9366 | 0.944 | 0.9959 |
| **4** | 0.9710 | 0.9708 | 0.9973 | 0.9375 | 0.944 | 0.9962 |
| **5** | 0.9709 | 0.9710 | 0.9966 | 0.9387 | 0.9445 | 0.9963 |
| **Mean** | 0.9711 | 0.9712 | 0.9970 | 0.9380 | 0.9445 | 0.9963 |
| **±Std Dev** | ± 0.0012 | ± 0.0011 | ± 0.0003 | ± 0.0013 | ± 0.0006 | ± 0.0003 |

**Supplementary Table S3. Per-label performance of the multilabel domain classifier on the DS150 held-out test set.**

All metrics were computed on the held-out test set (n = 15,000).

| **Domain** | **ROC-AUC** | **PR-AUC** | **MCC** | **F1** |
|---|---|---|---|---|
| NBS | 0.999 | 0.990 | 0.952 | 0.955 |
| LRR | 0.999 | 0.994 | 0.952 | 0.959 |
| TIR | 0.990 | 0.903 | 0.902 | 0.903 |
| CC | 0.976 | 0.817 | 0.701 | 0.703 |
| Kinase | 0.998 | 0.997 | 0.958 | 0.972 |
| **Macro average** | **0.992** | **0.940** | **0.893** | **0.899** |

**Supplementary Table S4. Classification counts and percentages for DREM proteome-wide predictions.** Absolute protein counts and percentages for each confidence class across the five plant proteomes, with summary statistics (mean, standard deviation, minimum and maximum).

| **Species** | **Total proteins** | **HC (count)** | **HC (%)** | **Candidate (count)** | **Candidate (%)** | **Low Conf (count)** | **Low Conf (%)** | **Non-resistance-like (count)** | **Non-resistance-like (%)** |
|---|---|---|---|---|---|---|---|---|---|
| *Arabidopsis thaliana* | 48,306 | 802 | 1.66 | 280 | 0.58 | 4,155 | 8.60 | 43,069 | 89.16 |
| *Oryza sativa* | 42,579 | 506 | 1.19 | 244 | 0.57 | 3,257 | 7.65 | 38,572 | 90.59 |
| *Solanum lycopersicum* | 34,361 | 378 | 1.10 | 234 | 0.68 | 2,687 | 7.82 | 31,062 | 90.40 |
| *Vitis vinifera* | 41,097 | 881 | 2.14 | 324 | 0.79 | 3,557 | 8.66 | 36,335 | 88.41 |
| *Cucurbita pepo* | 27,862 | 275 | 0.99 | 122 | 0.44 | 2,252 | 8.08 | 25,213 | 90.49 |
| **MEAN** | | 568.4 | 1.42 | 240.8 | 0.61 | 3,182 | 8.16 | 34,850 | 89.81 |
| **Std Dev** | | 263.9 | 0.48 | 75.2 | 0.13 | 742.1 | 0.45 | 6,907 | 0.97 |
| **Min** | | 275 | 0.99 | 122.0 | 0.44 | 2,252 | 7.65 | 25,213 | 88.41 |
| **Max** | | 881 | 2.14 | 324.0 | 0.79 | 4,155 | 8.66 | 43,069 | 90.59 |

**Supplementary Table S5. Classification, HMMER validation and resistance-protein classes across five plant proteomes.** Comprehensive DREM classification and validation results across five plant proteomes. The table reports absolute protein counts for the four DREM confidence classes; HMMER-confirmed and HMMER-divergent predictions; the distribution of resistance-protein classes within the High Confidence and Candidate tiers; and summary statistics for the binary resistance score. R-score values represent the resistance probability assigned by the binary classifier, whereas Proteins above threshold indicates the number of proteins exceeding the classification threshold.

| Metrics | Species | | | | |
|---|---|---|---|---|---|
| | *Arabidopsis thaliana* | *Oryza sativa* | *Solanum lycopersicum* | *Vitis vinifera* | *Cucurbita pepo* |
| **High Confidence, total** | 802 | 506 | 378 | 881 | 275 |
| **Candidate, total** | 280 | 244 | 234 | 324 | 122 |
| **Low Confidence/ Divergent** | 4,155 | 3,257 | 2,687 | 3,557 | 2,252 |
| **Non-resistance-like** | 43,069 | 38,572 | 31,062 | 36,335 | 25,213 |
| **HMMER-confirmed** | 811 | 585 | 450 | 939 | 269 |
| **HMMER-divergent** | 271 | 165 | 162 | 266 | 128 |
| **High Confidence TNL** | 256 | 1 | 24 | 38 | 1 |
| **High Confidence CNL** | 96 | 145 | 84 | 293 | 24 |
| **High Confidence NLR** | 85 | 112 | 52 | 198 | 9 |
| **High Confidence RLK/RLP** | 365 | 248 | 218 | 352 | 241 |
| **Candidate TNL** | 1 | 1 | 0 | 0 | 0 |
| **Candidate CNL** | 11 | 9 | 3 | 8 | 6 |
| **Candidate NLR** | 267 | 232 | 231 | 316 | 116 |
| **Candidate RLK/RLP** | 1 | 2 | 0 | 0 | 0 |
| **Mean R score** | 0.6313 | 0.5003 | 0.5956 | 0.6258 | 0.5849 |
| **Median R score** | 0.6254 | 0.4749 | 0.5689 | 0.6177 | 0.5691 |
| **Proteins above threshold** | 29,053 | 20,428 | 19,208 | 24,475 | 15,459 |

**Supplementary Table S6. HMMER validation of High Confidence and Candidate proteins across five plant proteomes, related to Table 1.** Proteins were classified as HMMER-confirmed when at least one significant domain hit (E-value < $1 \times 10^{-5}$) was detected for any of the five target Pfam profiles (NB-ARC, TIR, kinase, LRR and CC). HMMER-divergent proteins carry a DREM resistance signal but no detectable canonical domain.

| **Species** | **High Confidence + Candidate** | **HMMER-confirmed** | **HMMER-confirmed (%)** | **HMMER-divergent** |
|---|---|---|---|---|
| *Arabidopsis thaliana* | 1,082 | 811 | 74.95 | 271 |
| *Oryza sativa* | 750 | 585 | 78.00 | 165 |
| *Solanum lycopersicum* | 612 | 450 | 73.53 | 162 |
| *Vitis vinifera* | 1,205 | 939 | 77.93 | 266 |
| *Cucurbita pepo* | 397 | 269 | 67.76 | 128 |
| **Total** | **4,046** | **3,054** | **75.48** | **992** |

**Supplementary Table S7. Comparison of DREM and PRGminer, related to Fig. 5A**. Metrics on the held-out test set (7,542 resistance and 7,458 non-resistance proteins) were computed at a decision threshold of 0.5; values in brackets are 95% confidence intervals from 1,000 bootstrap resamples. For PRGminer, trainable parameters refer to the Phase 1 model and model size includes both phases. Throughput was measured on 2,000 sequences with the same GPU (NVIDIA Tesla T4).

| Metric | DREM | PRGminer |
|---|---|---|
| ***Held-out test set (n = 15,000)*** | | |
| Accuracy | 0.969 [0.966–0.972] | 0.910 [0.905–0.915] |
| Precision | 0.974 [0.971–0.978] | 0.897 [0.891–0.904] |
| Recall | 0.964 [0.960–0.968] | 0.927 [0.922–0.933] |
| F1-score | 0.969 [0.966–0.972] | 0.912 [0.907–0.916] |
| MCC | 0.938 [0.933–0.944] | 0.820 [0.811–0.829] |
| ROC-AUC | 0.997 [0.996–0.997] | 0.951 [0.947–0.954] |
| PR-AUC | 0.997 [0.997–0.998] | 0.938 [0.933–0.944] |
| Brier score | 0.021 [0.020–0.023] | 0.079 [0.075–0.082] |
| ***RefPlantNLR (n = 481)*** | | |
| Recall | 478/481 (99.4%) | 478/481 (99.4%) |
| ***Efficiency*** | | |
| Trainable parameters | 183,457 | $50.7 \times 10^{6}$ |
| Model size | 0.74 MB | 613 MB |
| Throughput (sequences/s) | 2,829 | 80 |

McNemar test on the held-out test set: 13,404 proteins correctly classified by both tools, 1,133 only by DREM, 245 only by PRGminer and 218 by neither ($\chi^2$ with continuity correction, $p = 3.5 \times 10^{-126}$). MCC, Matthews correlation coefficient; PR-AUC, area under the precision–recall curve; ROC-AUC, area under the receiver operating characteristic curve.
